# Flies tune the sensitivity of their multifunctional gyroscope

**DOI:** 10.1101/2024.03.13.583703

**Authors:** Anna Verbe, Kristianna M. Lea, Jessica L. Fox, Bradley H. Dickerson

## Abstract

Locomotion requires navigating unpredictable and complex environments, demanding both stability and maneuverability within short timeframes. This is particularly important for flying insects, and the true flies (Diptera) stand out among this group for their impressive flight capabilities. Flies’ aerial abilities are partially attributed to halteres, tiny club-shaped structures that evolved from the hindwings and play a crucial role in flight control. Halteres oscillate during flight, in antiphase with the wings, providing rhythmic input to the wing steering system *via* arrays of embedded mechanosensors called campaniform sensilla. These sensor arrays convey timing information to the wing steering muscles, but linking haltere sensor location to sensor activity and the functional organization of the wing steering system remains a central challenge. Here, we use *in vivo* calcium imaging during tethered flight to obtain population-level recordings of the haltere sensory afferents in specific fields of sensilla. We find that haltere feedback is continuously modulated by visual stimuli to stabilize flight. Additionally, this feedback is present during saccades and help flies actively maneuver. We also find that the haltere’s multifaceted role arises from the steering muscles of the haltere itself, regulating haltere stroke amplitude to modulate campaniform activity. Taken together, our results underscore the crucial role of biomechanics in regulating the dynamic range of sensors and provide insight into how the sensory and motor systems of flies coevolved.

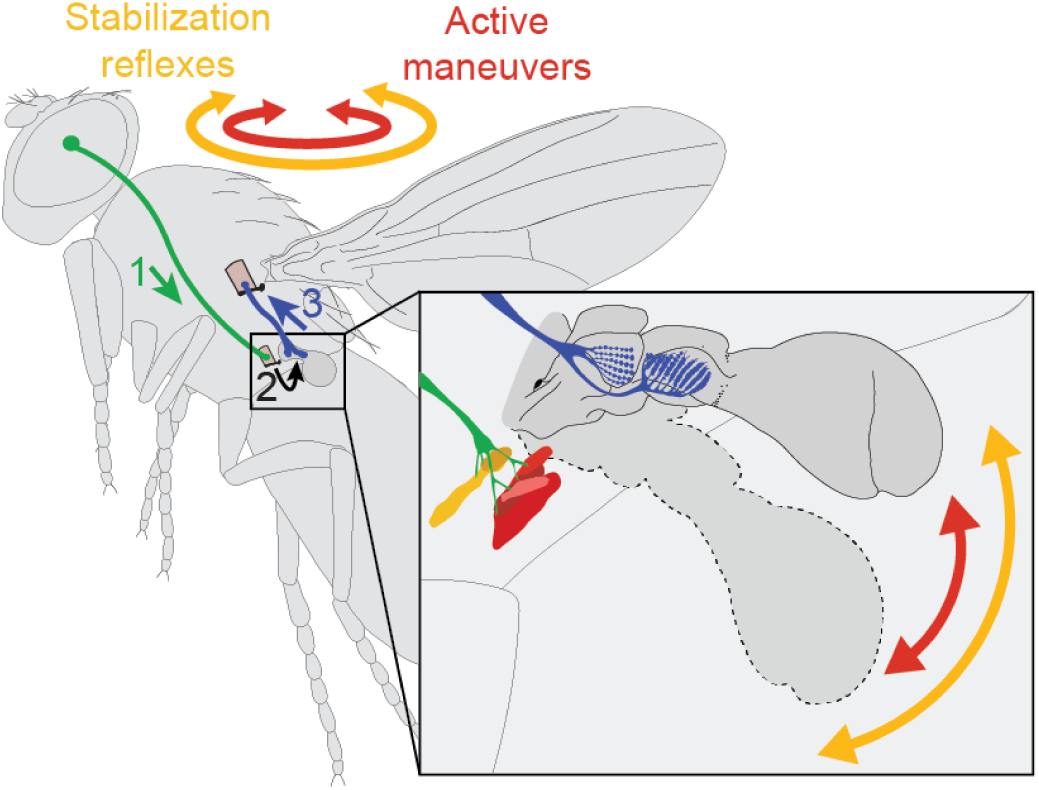

## Introduction

Animals navigate unpredictable and challenging three-dimensional environments. This requires the ability to both remain stable, to counteract perturbations, and be maneuverable, to actively pursue prey or escape a predator at short timescales. As a result, neural circuits that detect precise spike timing are essential to survival, and many organisms use specialized organs to detect tiny timing differences^1–7^. Organs such as the ears of barn owls or the electric organ in knifefish have large arrays of mechanosensory cells that possess specific anatomical and mechanical properties that biomechanically filter the kinetic energy of a stimulus, thereby extracting relevant information while minimizing neural processing time^8–11^. Although several circuits that detect timing differences have been identified, how the encoding by specialized sensors relates to the functional organization of the motor system is less understood. This is a critical knowledge gap as the timing of neural input has important consequences on the biomechanical properties of the musculoskeletal system^12–17^, which ultimately executes behavior.

These issues are particularly pronounced for flying insects, which collect and process incoming information to control wing motion at sub-millisecond timescales^18^. Among flying insects, true flies (order Diptera) stand out for their adept aerial maneuvers. Indeed, flies have evolved multiple physiological specializations in both the sensory and motor systems that make them remarkable fliers and one of the most diverse insect orders^19^.

For example, flies possess a unique neural superposition visual system that helps process optic flow at high speeds^20^. Furthermore, in flies (as in many other insect orders) the flight muscles, which reside solely in the thorax, are functionally segregated into two major groups: power muscles that provide the force necessary to flap the wings and generate lift, and steering muscles that control the subtle changes in wing motion needed to accomplish maneuvers^21,22^. Exoskeletal deformation during flapping activates the power muscles *via* stretch to achieve the high frequencies needed to produce lift. These muscles are ‘asynchronous:’ a single action potential triggers multiple contraction cycles^23^. By contrast, the wing steering muscles, each innervated by a single motor neuron, are ‘synchronous’; their firing times rely on mechanosensory feedback that arrives during each wingstroke. There are two major sources of mechanosensory feedback that structure the firing times of the steering muscles: the wings themselves, and vestigial hindwings unique to flies called halteres^24–26^.

Halteres are tiny club-shaped structures found on the metathorax that do not serve an aerodynamic function but instead provide sensory information crucial to flight^27–30^. Halteres oscillate during flight, providing rhythmic input to the wing steering system *via* arrays of embedded mechanosensors (campaniform sensilla) that are sensitive to cuticular strain^31^. Both the wings and halteres possess campaniform sensilla, but they have hypertrophied on the haltere such that in the case of *Drosophila melanogaster*, the haltere has approximately 140 campaniforms compared to less than 50 on the wing^32^.

The haltere is also the only true biological gyroscope: during rotations, it experiences Coriolis forces due to its tendency to resist changes its plane of oscillation, triggering equilibrium reflexes of the head and wings^28–30,33–37^. Underscoring their evolutionary history as a hindwing, halteres are equipped with a single power muscle and a set of seven control muscles that are serially homologous to those of the wings^38–42^. These muscles receive visual input, allowing flies to modulate the activity of the wing steering system *via* the halteres even in the absence of body rotations, a process known as the control-loop hypothesis^22,39,42^ (Fig. 1A). Through active control of the haltere muscles, flies may execute voluntary movements without triggering counteracting reflexes. Thus, halteres serve as multifunctional sensory organs that provide essential timing information to flight circuitry. However, how this multifunctional role is achieved is poorly understood.

**Figure 1:**
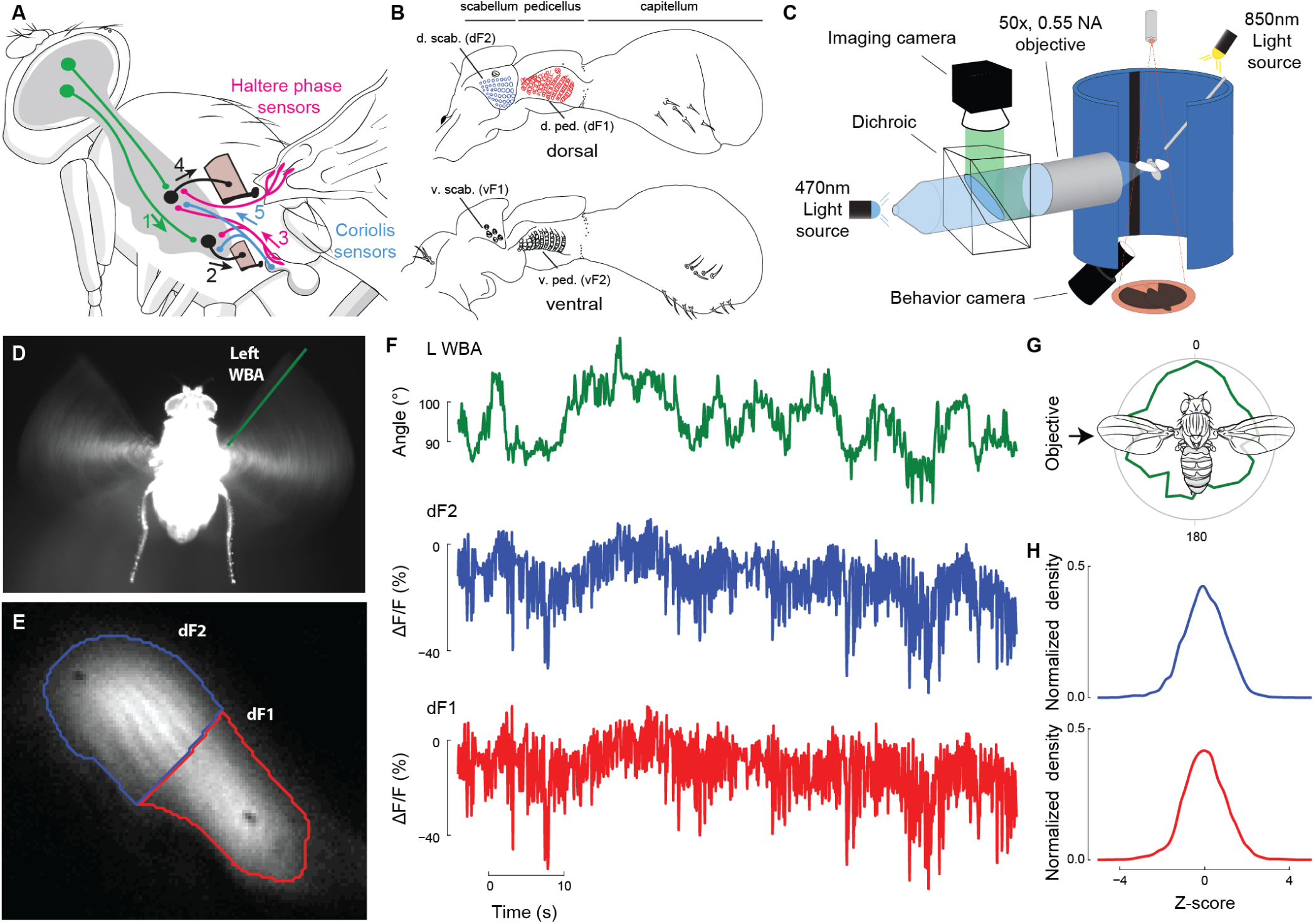
*In vivo* imaging of the dorsal haltere campaniform fields during tethered flight. (A) The control-loop hypothesis explains the multi-sensory activity of the halteres (redrawn from Dickerson *et al*.^42^). Visual signals (1) are sent to the haltere muscles (2), recruiting additional campaniform sensilla with different preferred firing times (3). This feedback alters the timing or activation of the wing steering muscles (4). The haltere’s gyroscopic function may operate through a similar pathway (5). (B) Anatomy of the *Drosophila* haltere, showing the locations of the four major campaniform fields (redrawn from Cole and Palka^32^). (C) Schematic of setup used to image activity of the dorsal campaniform fields on the left haltere while simultaneously tracking wing motion during the presentation of visual stimuli. (D) Image from the behavioral tracking camera showing the fly from below and the wingstroke envelope of both wings; left wing (ipsilateral) amplitude is shown in green line. (E) Maximum intensity projection of the dorsal campaniform fields of a single experiment, clustered by fluorescence activity into regions that correspond with dF1 (red) and dF2 (blue). (F) Extracted signals of the left wingbeat amplitude, fluorescence of the mean area of dF2 and dF1. (G) Polar probability density histogram showing the distribution of stripe azimuth during closed-loop fixation behavior. (H) Mean histograms of fluorescence changes during stripe fixation, normalized as z-scores. N =23 flies. See Supplemental Movie 1.

Across flies, the campaniform sensilla embedded in the haltere are arranged in highly stereotyped groups, called fields^28,32,43–46^ (Fig. 1B). Each field has a different spatial orientation that may relate to its activity patterns in flight^28,44^. Although longstanding hypotheses suggest that these different arrays may provide different information relevant to flight control^28,44^, we know little about how haltere sensor location maps to sensory activity and how this activity, in turn, relates to the functional organization of the wing steering system. We leveraged the genetic tools available in *Drosophila* to investigate how visual input modulates the encoding of two campaniform fields, dF1 and dF2, by imaging neural activity from a moving haltere during tethered flight. We then probed how this encoding maps to the organization of the wing steering system and is fundamentally determined by the kinematics of the haltere. Our results demonstrate the crucial role of biomechanics in regulating the dynamic range of sensors so that they can mediate both stabilizing and active maneuvers.

## Results

### *In vivo* imaging of the haltere’s sensory fields

To image from the haltere campaniform fields during tethered flight, we used a driver line that target most of the population of haltere campaniform sensilla, *DB331-GAL4*, driving the expression of the genetically encoded calcium indicator GCaMP7f (Fig. 1). The neurons associated with each campaniform field are directly beneath the dome of each sensilla and the cuticle in this region is quite thin. This allowed us to image GCaMP activity with an epifluorescent microscope using a previously described method^42,47^ in an intact animal (Fig. 1C). We simultaneously measured wing steering effort, reported here as left-minus-right wingbeat amplitude (L-R WBA), using custom machine vision software (Fig. 1D, Supplemental Movie 1). Due to the high density of campaniforms within each field, we could not resolve GCaMP activity at the level of individual sensilla. We instead report the activity of each field from the left haltere using k-means clustering of the image stream to extract signals for the dorsal campaniforms fields, dF1 and dF2 (Fig. 1E). Campaniforms fire single, phase-locked action potentials in response to periodic, oscillatory stimuli^48,49^. Additionally, increasing wing bending recruits campaniform firing at different preferred stimulus phases^48^. We therefore interpret increases in GCaMP signal as recruitment of additional sensilla. Preliminary experiments indicated that the ventral fields are also active in flight. However, these are subject to substantial motion artifacts, so we focused on the calcium activity of the dorsal fields.

Under closed-loop conditions where the fly controls the position of a dark stripe in its visual field *via* changes in L-R WBA, haltere feedback from both dF1 and dF2 is modulated throughout flight (Fig. 1F, G). The patterns of activity from dF1 and dF2 are correlated with wingbeat amplitude activity over time. In control experiments GFP expression in the haltere campaniforms demonstrated that these signal fluctuations were not motion artifacts. Pooling data across flies allowed us to test whether these modulations were consistent. We normalized each fluorescence trace, creating z-scored versions of the data from all 23 flies, and subsequently created population histograms of dF1 and dF2 activity during closed-loop stripe fixation (Fig. 1H). The fluorescence activity of both dF1 and dF2 were normally distributed, indicating that the activity of each field fluctuated about some baseline level.

### Haltere feedback is directionally tuned

We next tested the sensitivity of the dorsal haltere campaniform sensilla to an array of open-loop visual stimuli. Previous calcium imaging of the haltere axon terminals demonstrated that their activity is modulated by rotations of wide-field visual motion^42^. We subjected flies to simulated rotations in the sagittal (yaw-roll) and azimuthal (pitch-roll) planes while simultaneously tracking changes in wingstroke amplitude. Using random starfield patterns, we shifted the center of rotation in 30° increments and compared the kinematic changes to the patterns of dF1 and dF2 activity elicited by the corresponding patterns of visual motion (Fig. 2A, Fig. S1).

**Figure 2:**
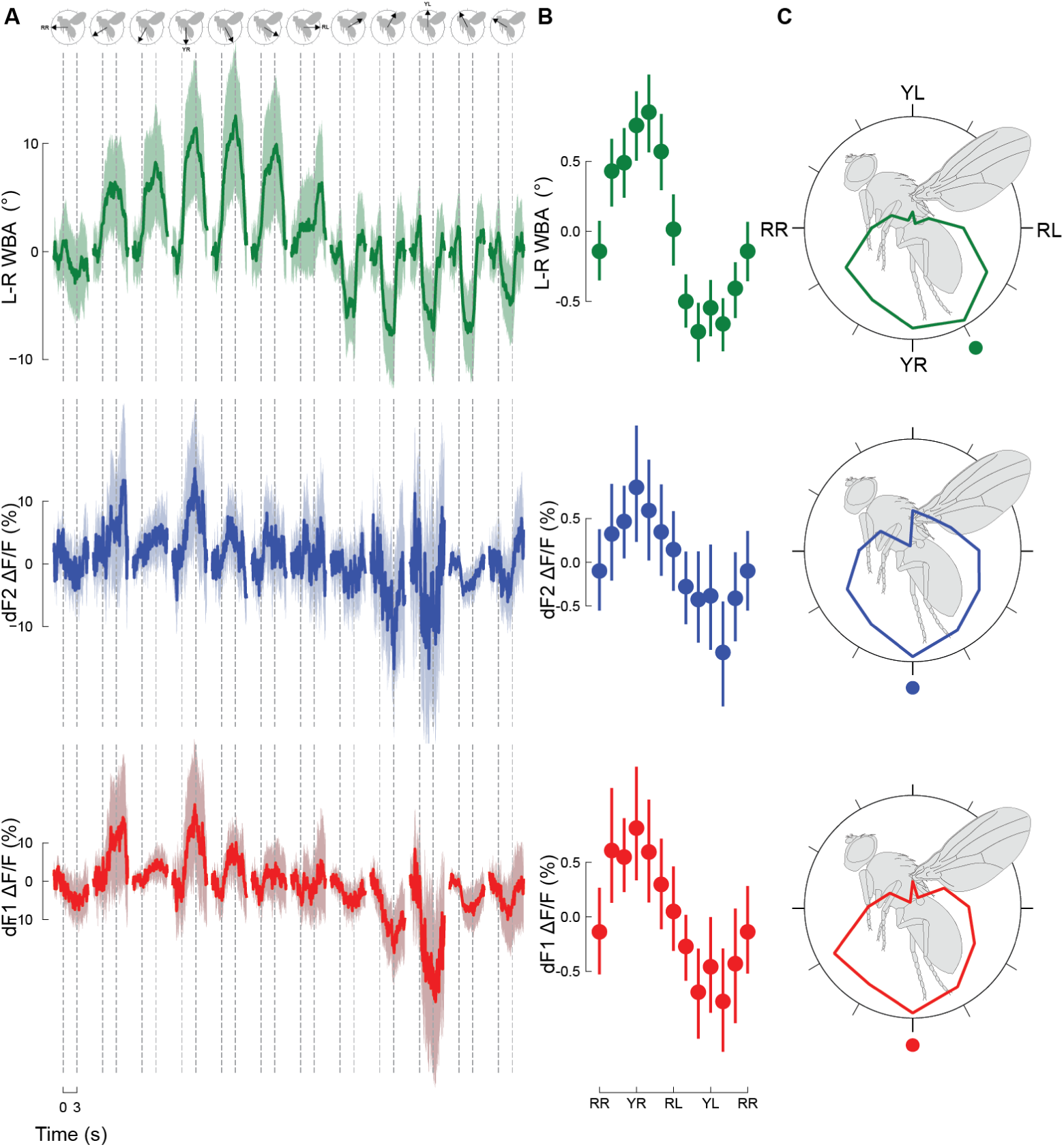
Haltere sensory feedback is directionally tuned by visual motion. (A) Tuning dynamics of the dorsal campaniform sensilla fields in response to 3 s presentations of wide-field visual rotational motion about the yaw-roll axis. Stimulus direction (top), wingbeat amplitude responses, and campaniform fields fluorescence where the stimulus center of rotation shifted in 30° increments. Data represent means ±95% confidence intervals (CIs). RR, roll left; RR, roll right; YL, yaw left; YR, yaw right. Roll right is plotted twice to emphasize the cyclical nature of the data. Stimulus direction is indicated by the right-hand rule. (B) Stroke amplitude and haltere campaniform field activity plotted as functions of rotational orientation in the midsagittal plane. The curves were constructed by determining the average signal over the 3 s following stimulus onset. (C) Mean tuning functions from (B) plotted in polar coordinates. Solid dots indicate the orientation of the maximum response for each signal. n=21 flies. See also Supplemental Movie 1.

From these data, we calculated tuning curves for L-R WBA as well as dF1 and dF2 activity (Fig. 2B, C). For presentations of wide-field motion about the yaw-roll axis, L-R WBA and the calcium activity of both dF1 and dF2 varied sinusoidally with the stimulus rotation angle, with a maximum response near yaw motion to the right and a minimum response around yaw to the left (Figure 2B, C). Notably, the tuning of dF1 and dF2 matches the sensitivity of flies’ steering effort, yet the campaniform fields demonstrate tuning in the opposite direction of the haltere steering muscles^42^. Although visual rotations about the pitch-roll axis elicited changes in wing kinematics, neither dF1 nor dF2 show significant changes in calcium activity or directional selectivity (Fig. S1).

### Haltere feedback is linearly recruited for visual stabilization reflexes and nonlinearly recruited for active maneuvers

We then examined the relationship between haltere campaniform activity and the functional organization of the wing steering system. The wing steering muscles are functionally subdivided into two classes, tonically active muscles that are linearly recruited for stabilization reflexes, and phasically active muscles that have a steep nonlinear activation threshold and are recruited for active maneuvers^47,50–52^.

We first asked how visual stimuli shape the relationship between the activity of the campaniform fields and steering effort magnitude. If the halteres help regulate the activity of the tonically active muscles, then modulations of haltere campaniform field activity and steering maneuver magnitude should be linearly related. To test this, we sorted the L-R WBA data from the responses to wide-field visual motion, regardless of the direction of stimulus motion, along with the corresponding calcium signals for dF1 and dF2. We sorted 1260 responses from 21 flies from the most leftward to the most rightward responses during the 3 s following stimulus onset (Fig. 3A). We then divided the sorted data into amplitude deciles ranging from the largest leftward responses (90^th^–100^th^ percentile) to the largest rightward responses (0^th^–10^th^ percentile). We then calculated the mean signal for each decile during the stimulus (Fig. 3B). The average peak haltere responses during the stimulus windows in each stroke amplitude decile are plotted in Figure 3C. For both dF1 and dF2, we observed similar patterns of recruitment as a function of L-R WBA (Fig. 3B, C). Although the behavioral and campaniform field signals are individually nonlinear (Fig.3), there is a linear relationship between L-R WBA and either dF1 or dF2 (Fig. 3D).

**Figure 3:**
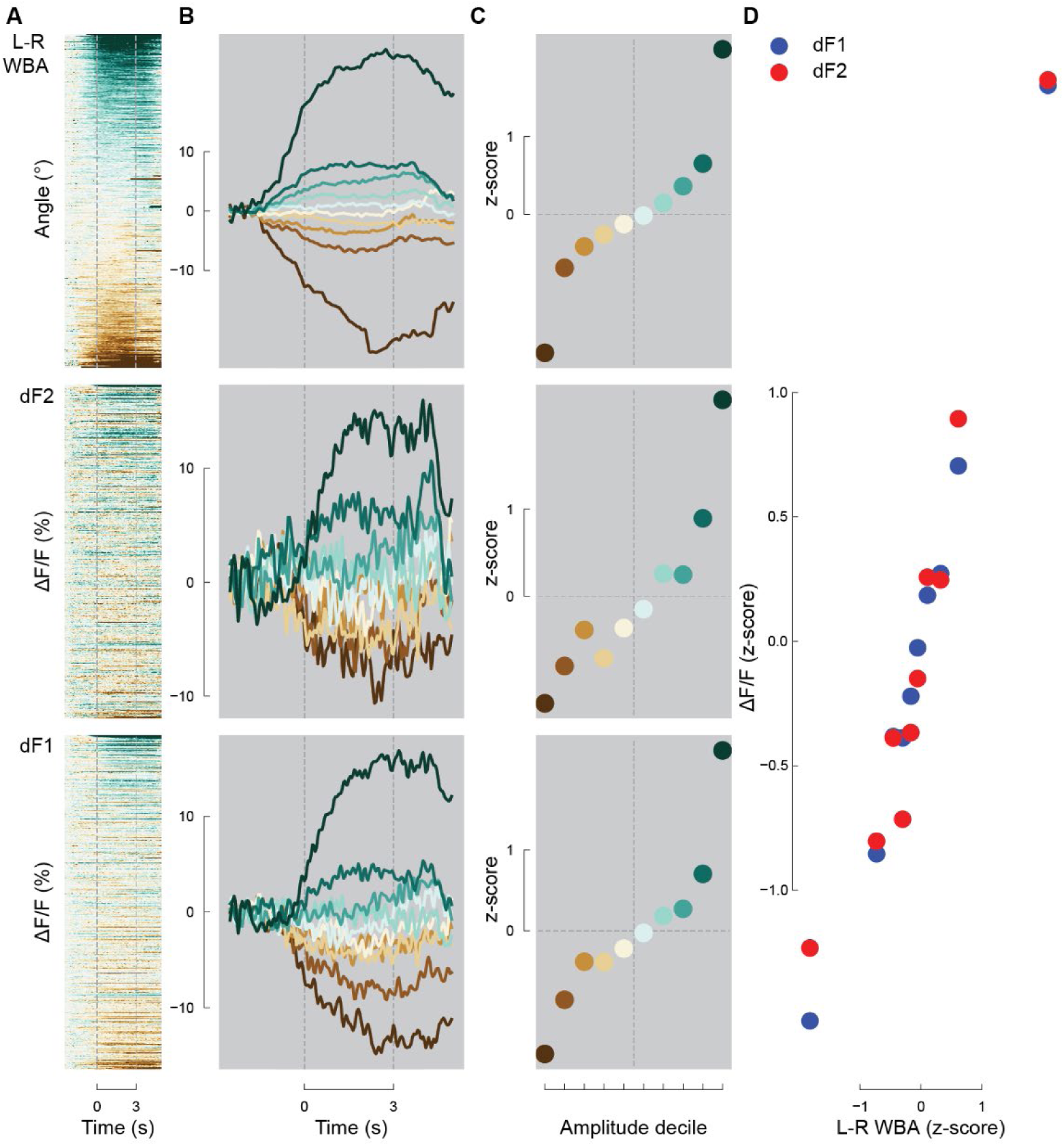
Haltere feedback is linearly recruited for visual stabilization reflexes. (A) Raster plots showing the left-minus-right wingbeat amplitude and the corresponding fluorescence signals of dF1 and dF2 in response to a 3 s episode of wide-field motion about the yaw-roll axis. Data are sorted in descending order according to the average magnitude of ipsilateral stroke amplitude during the 3 s following stimulus onset. (B) Decile mean responses of wing kinematics and campaniform field fluorescence, ranging from the largest rightward turns (0^th^–10^th^ percentile, green) to the largest leftward turns (90^th^–100^th^ percentile, brown). (C) Plots showing mean response in the 3 s episode for each decile. (D) Mean decile response of dF1 and dF2 calcium signals during 3 s stimulus presentation compared to L-R WBA. n=21 flies.

If the haltere is involved in active flight maneuvers, campaniform field activity will be nonlinearly related to turn magnitude. We therefore next asked how haltere feedback may be modulated during active maneuvers. Flies exhibit rapid redirection maneuvers during flight named saccades. Although most saccades are triggered by external visual stimuli^53,54^, flies also exhibit spontaneous saccades^55^, and they are conspicuous features of tethered flight preparations^56^ and haltere gyroscopic input terminates saccades^57^. We used a simple classifier^58^ (based on L-R WBA) to identify saccades and the corresponding calcium signals of dF1 and dF2. We applied our classifier to a total of 4,521 saccades from 23 flies that we imaged during closed-loop stripe fixation (Fig. 4A). We again sorted these data into amplitude deciles based on L-R WBA magnitude and calculated decile means (Fig. 4B). In contrast to the linear recruitment of haltere feedback during wide-field visual motion (Fig 3D), we observed a nonlinear relationship between steering effort and campaniform field activity (Fig 4D). Whereas L-R WBA monotonically increased, we find that dF1 and dF2 are either inactive or show low activity for the bottom 50% of identified saccades, and linearly recruited for the remaining 50% (Fig. 4C). Notably, these modulations in dF1 and dF2 activity occur before the peak changes in L-R WBA during a saccade. We also looked at spontaneous saccades during presentation of a static visual pattern but saw no consistent trends in dF1 or dF2 activity (Fig. S2). Together, these results indicate that dF1 and dF2 are recruited in a linear fashion during visually mediated reflexes (Fig. 3D) and a nonlinear fashion prior to executing saccades (Fig. 4D).

**Figure 4:**
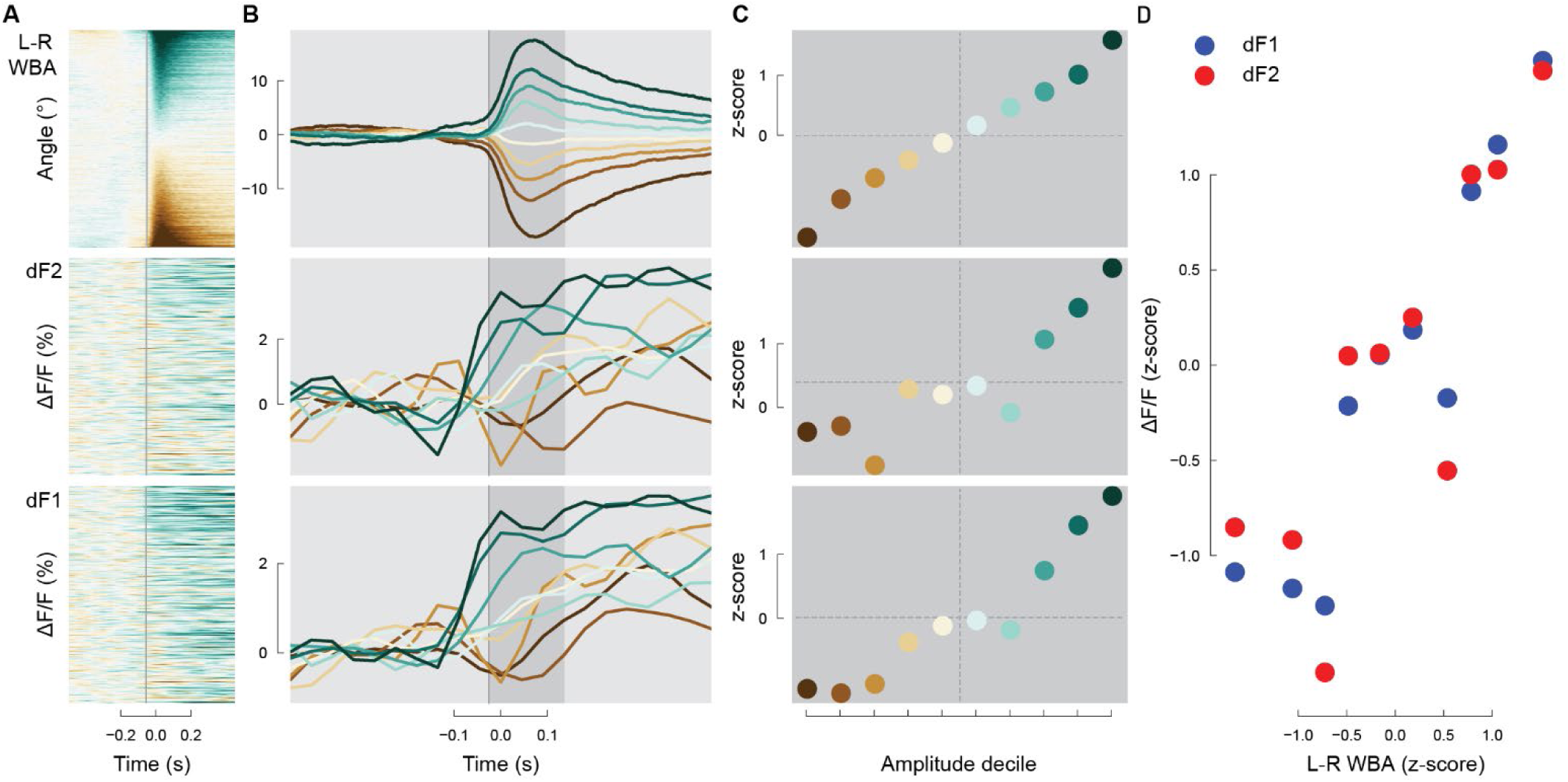
Haltere feedback is nonlinearly recruited for active maneuvers. (A) Raster plots of all saccades sorted by the change in left-minus-right wingbeat amplitude. (B) Saccade-triggered averages of wing kinematics and dorsal halteres signals sorted into amplitude deciles, ranging from the largest rightward turns to the largest leftward turns. Dark gray bar denotes saccade initiation. (C) Plots showing mean response for each decile during the saccade. (D) Mean decile response of dF1 and dF2 calcium signals during 3 s stimulus presentation compared to L-R WBA. n = 4521 saccades from 23 flies.

### The haltere muscles are differentially recruited

Modulation of haltere feedback during both visually mediated stabilization reflexes and active maneuvers can be explained by two scenarios. Either the campaniforms embedded within each field exhibit context-dependent encoding or the haltere motor system is functionally stratified. Although campaniforms exhibit either rapid or slow adaptation to applied static loads^49,59^, when stimulated at wingbeat frequency, they all fire single spikes at different preferred phases^48,49,60^. This suggests that the haltere steering muscles are functionally segregated like their forewing counterparts.

To test the hypothesis, we expressed GCaMP6f in a driver line (*R22H05-GAL4*) that targets all the haltere steering muscles. The haltere steering muscles are divided into two major anatomical groups: the basalares (hB1 and hB2) and the axillaries (hI1, hI2, hIII1, hIII2, and hIII3; Fig. 5A)^42^. We imaged haltere muscle activity using the same method we applied to the haltere campaniform fields (Fig. 5B) and again identified spontaneous saccades. We classified a total of 2,627 saccades from 7 flies during the presentation of a closed-loop visual pattern. From the identified saccades, we sorted the calcium signals associated with each muscle group into deciles and again calculated averages for each group (Fig. 5C). The activity of the haltere basalares is linearly related to saccade amplitude, consistent with tonically active wing steering muscles (Fig. 5D)^47^. The activity of the haltere axillaries, however, is nonlinearly related to saccade amplitude, consistent with phasically active muscles (Fig. 5D). We interpret the recruitment of the axillaries to represent separate muscles, each of which are recruited during large turns of opposite directions. Thus, the haltere motor system is both anatomically and functionally stratified.

**Figure 5:**
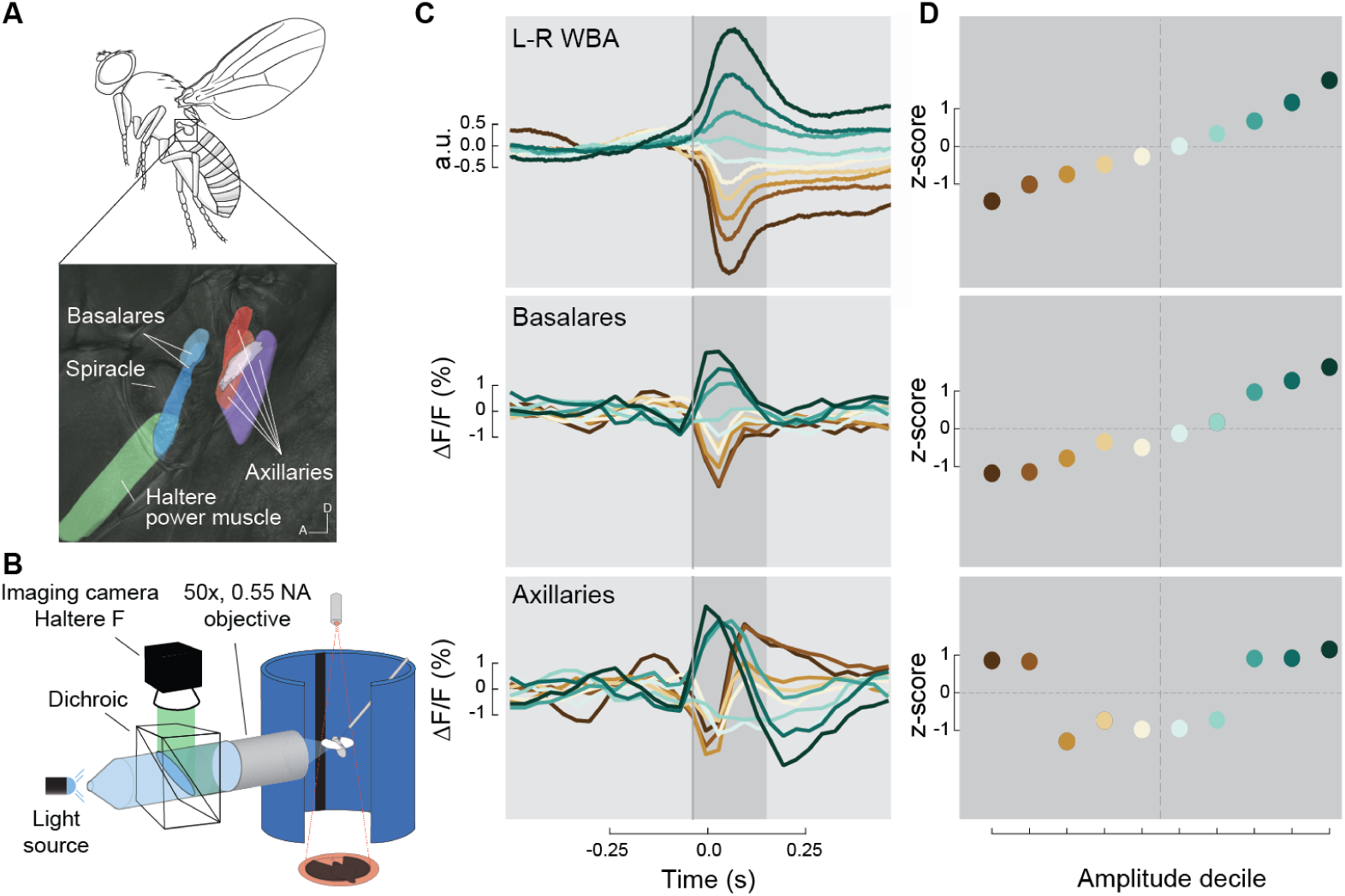
The haltere muscles are differentially recruited. (A) The haltere motor system of *Drosophila* consists of a power muscle and seven steering muscles that can be divided into two anatomical groups, the basalares and the axillaries. (B) Schematic of setup used to simultaneously image muscle activity and track wing motion in response to visual stimuli. A and B are reproduced from Dickerson *et al.*^42^ (C) Saccade-triggered averages of left-minus-right wingbeat amplitude and haltere muscle fluorescence for each anatomical group. Saccade-triggered averages are plotted as means of each amplitude decile, sorted from the largest rightward turns to the largest leftward turns. (D) Plots showing the mean decile response during a saccade (dark grey bar in C). n = 2,627 saccades from 7 flies.

### Optogenetic activation of haltere steering muscles changes haltere amplitude

Changes in haltere campaniform field activity result from the activity of the haltere steering muscles, which regulate either kinematics or the biomechanics of each campaniform field^22,39,42^. Although these two modes of regulating haltere sensing are not mutually exclusive, wing campaniform activity is correlated with changes in L-R WBA in response to wide-field visual motion stimuli, much like haltere campaniforms^42^. Changes in wing kinematics, and the resulting strain on the wing, are the direct result of wing steering muscle activity. Given their serial homology with the wings, this suggests that the haltere muscles regulate stroke amplitude to modulate campaniform recruitment and, ultimately, wing steering muscle activity. To test if the haltere steering muscles control haltere kinematics, we optogenetically activated two haltere muscle motor neurons during tethered flight and recorded haltere motion at 2,000 fps using a high-speed camera (Fig. 6A). We expressed Chrimson in a driver line (*SS41075-SplitGAL4*^61^) that targets two haltere muscle motor neurons: hDVM, the asynchronous power muscle^62^, and hI1, an axillary steering muscle (Fig. 6B). Optogenetic activation of these motor neurons decreased haltere stroke amplitude (Fig. 6C-F, Supplemental Movie 2). The reduction in amplitude is a result of the haltere’s failure to reach its full ventral extent during the stimulus; the dorsal extreme is not as strongly affected.

**Figure 6:**
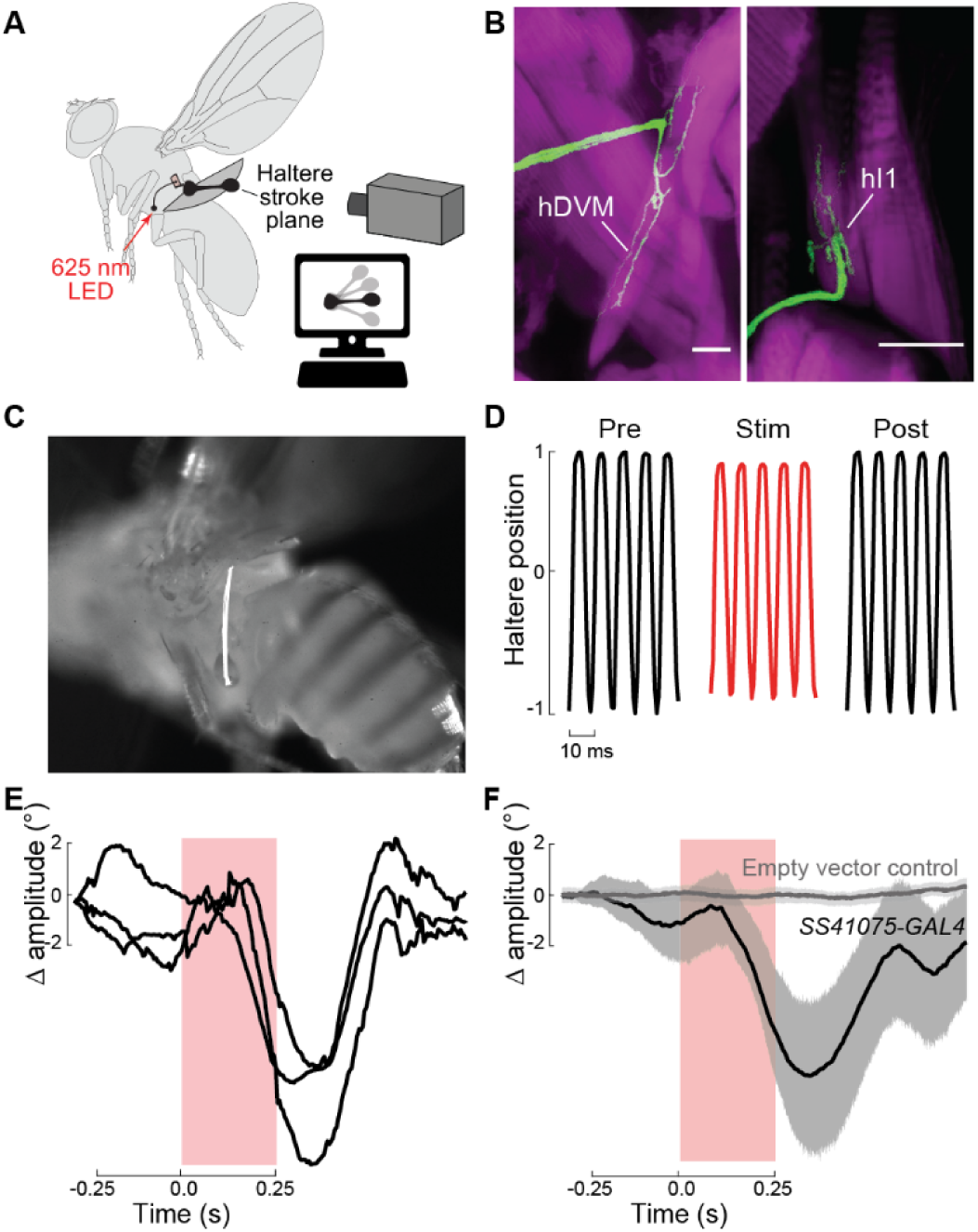
Optogenetic activation of haltere steering muscle motor neurons decreases haltere stroke amplitude. (A) Schematic of setup used to record haltere motion of tethered, flying flies during optogenetic activation of haltere motor neurons. (B) Maximum intensity projections of the haltere power muscle, hDVM, and one steering muscle motor neuron, hI1, expressing GFP (green) driven by *SS41075-GAL4*. Magenta shows phalloidin staining of muscles. Images reproduced from Dickerson *et al*.^42^ (C) Sample frame of haltere tracking with the trajectories of 10 tracked strokes overlaid (white). (D) Example traces of normalized haltere stroke amplitude 25 ms before or after (black) and during optogenetic activation (red). (E and F) Three representative traces (E) and population data (F) showing the change in haltere stroke amplitude following optogenetic activation of *SS41075-GAL4*. Changes in haltere stroke amplitude for empty vector control flies are also plotted on the right (grey). Data shown represent mean ± 95% CI, n =54 flies. See also Supplemental Movie 2.

## Discussion

Whereas previous electrophysiological work demonstrated how haltere neurons encode mechanical input using quiescent flies^25,26,28,49,63,64^, here we used *in vivo* calcium imaging during tethered flight to obtain population-level recordings of the haltere afferents in specific fields of sensilla. We provide further evidence that haltere feedback is continuously modulated by visual stimuli to stabilize flight and we also found that such feedback may trigger saccades and help flies actively maneuver. These different roles for flight control are determined by the haltere motor system, which subtly alters haltere kinematics to modulate campaniform activity. Thus, our results demonstrate how biomechanics can filter sensory stimuli to help produce flexible and robust behavior.

### Role of haltere in visually mediated flight

Like other insect campaniform sensilla, the orientation and location of the haltere fields are hypothesized to have functional significance^28,44,65,65–68^. For example, the fields of sensilla that lie parallel to the haltere’s long axis–i.e., dF1, vF1, and vF2–are predicted to detect the in-plane strains produced by the large vertical oscillations^28,44^. Similarly, the diagonally orientated sensilla of dF2 should be maximally sensitive to the strains produced either by lateral bending from gyroscopic torques or from active lateral haltere movements from the activity of the haltere steering muscles^28,39,42,44^. We therefore expected to observe that dF1 alone is continuously active during tethered flight, and that df2 is recruited during active maneuvers.

Instead, we found that the activity of both dF1 and dF2 are continuously regulated during flight (Fig. 1). These results are significant in two respects. First, the observation that dF2 activity is modulated by specific directions of wide-field visual motion (Fig. 2) provides further confirmation of the control-loop hypothesis^39,42^. The tuning characteristics of dF2, and its alignment with steering responses, suggest that the haltere steering muscles manipulate haltere motion to modulate haltere feedback. Indeed, our experiments provide the first direct evidence that the haltere steering muscles regulate stroke amplitude during flight (Fig. 6). This finding is consistent with previous work showing that the haltere steering system is also tuned to visual motion, yet its peak sensitivity is in the opposite direction of wing steering responses^42^. Thus, the haltere motor system may regulate campaniform field activity *via* tiny changes in flapping amplitude that depend on the wide-field visual environment.

Second, our results suggest that dF1 may mediate flight control. Previous anatomical and physiological work in the blowfly *Calliphora* showed that haltere afferents provide excitatory input to the wing motor neuron b1, through a mixed electrotonic and chemical synapse^25,26,69^. However, this feedback is provided by a single field of campaniforms, dF2: ablating dF2 and stimulating the remaining campaniform fields, including dF1, failed to generate postsynaptic potentials in the b1 motor neuron^26^. Yet in our experiments, dF1 is recruited in much the same manner as dF2, for both visually mediated reflexes and active maneuvers (Fig. 2-4), indicating that dF1 is not merely a passive sensor for the haltere’s large oscillation. Moreover, anatomical work shows extensive projection patterns of each field within the fly central nervous system^69^, suggesting that all fields play an important part in controlling flight. Future work with new connectomics data will reveal the organizational logic of the haltere campaniform fields with the wing steering motor neurons and other targets.

More broadly, the observation that both dorsal fields are continuously active is not surprising as the haltere undergoes large oscillations at a high frequency, creating complex patterns of strain across the surface. These strain patterns are transduced by the embedded campaniform sensilla, which are exquisitely sensitive to the slightest deformations of the cuticle^31,49,64,70^. Our use of a calcium indicator to monitor haltere activity allowed us to obtain population recordings from the two dorsal fields. However, the kinetics of our calcium indicator are too slow to capture crucial information about spike timing from individual sensilla. Intracellular recordings from sensilla axons demonstrate that changes in haltere motion are reported *via* either shifts in the preferred firing phase of active campaniforms or recruitment of additional campaniforms that have unique preferred firing phases^64^. By adjusting haltere stroke amplitude *via* modulation of the haltere steering muscles, flies may modulate the strength of this feedback by subtle changes in stroke amplitude to rapidly adjust wing kinematics with precisely timed mechanosensory feedback.

### Haltere feedback during active maneuvers

In addition to suggesting that visual information modifies haltere mechanosensory feedback and consequently wing motor output, the control-loop hypothesis predicts that the haltere control muscles and sensilla are active during voluntary maneuvers, e.g., body saccades. Consistent with their role in sensing the Coriolis forces that result from body rotations^57^, feedback from the halteres terminates saccades. However, whether haltere feedback is modulated during the initiation of saccades remains an open question^71^.

We found that both the haltere motor system, as well as the campaniform fields, are recruited during spontaneous saccades (Fig. 4, 5). Moreover, these changes in muscle and campaniform field activity occur prior to the changes in wing kinematics. This suggests that haltere motion can be controlled independently of the wings to initiate a voluntary maneuver. Recent work shows that flies tune the magnitude of a saccade according to the angular velocity of a visual stimulus^72^. Similarly, we found that flies can actively tune the strength of haltere mechanosensory feedback depending on the magnitude of a saccade (Fig. 4).

Recent work in insects suggests a role for efference copy in suppressing visually mediated reflexes during voluntary flight maneuvers^58,73,74^. For example, electrophysiological recordings of lobula plate tangential cells (LPTCs) in *Drosophila* show predictive scalable inhibition or excitation, correlated with spontaneous saccades^58,74^. Yet in all cases, the source of these efference copies remains unidentified. By contrast, our results suggest that rather than cancel out expected haltere feedback during a turn, flies co-opt existing haltere reflex loops to actively maneuver. An efference copy signal would render a fly susceptible to mechanical perturbations. Instead, by actively manipulating the haltere during a turn, flies can remain sensitive to gyroscopic forces during voluntary maneuvers.

### How does a fly reconcile internal mechanics with Coriolis force sensing in flight?

The multifunctional capability of the haltere system suggests that flies can be maneuverable at low cost to their stability^22^. However, how flies can distinguish self-motion mechanosensory feedback from external whole-body mechanical perturbations remains unclear. Campaniform sensilla function through a generic encoding mechanism, termed Derivative Pair Feature Detection (DPFD)^75^. DPFD may be an inherent property of neurons with non-specialized Hodgkin-Huxley dynamics, like many mechanosensors. As a result, the anatomical placement and mechanics of a DPFD neuron, rather than the specialized neural computation and membrane dynamics, act as a biomechanical filter and may confer specialized encoding. We hypothesize that the orientation of the different campaniform fields on the haltere allows for the control-loop and gyroscopic sensing to work synergistically to control flight maneuvers. In this scheme, descending visual commands to the haltere steering system can initiate a turn by causing small changes in haltere kinematics and by recruiting campaniforms from all fields. Then, as the fly begins to rotate and the haltere undergoes changes in its trajectory due to Coriolis forces, the activity of dF2–which should be maximally sensitive to the resulting shear strains–is increased, triggering a corrective maneuver. This hypothesis is consistent with both our observation that haltere feedback is constant and the haltere’s well-established role as a gyroscopic sensor.

In free flight, both stabilization and active maneuvers consist of banked turns that involve coordinated changes about all three cardinal axes^76,77^. Modeling and behavioral evidence show that the haltere can detect any combination of body rotations^30,34,36,78,79^. In this regard, it is surprising that visual modulation of campaniform sensilla activity seems to be only present in the yaw-roll plane and nonexistent for rotations about the pitch-roll plane. This is of particular interest as the direction of motion that elicited the strongest response, for both dF1 and dF2, is pure yaw, which is the weakest axis for gyroscopic responses^36^. It is possible that, in addition to helping initiate active turns, one major role for the control-loop is to help mediate straight flight about the azimuth by instituting rapid corrections. Indeed, flies dedicate a great deal of neural circuitry to descending interneurons that are hypothesized to maintain a straight flight trajectory^80^. Presumably some of those commands are directed to the haltere motor system.

### Co-evolution of the haltere and flight

Although the haltere is a unique sensory structure, its core function–regulating the timing of the wing steering system–and control are much like the aerodynamically functional forewings. Indeed, haltere evolved from the hindwing^19^, and past genetic work confirms that it is a serially homologous structure^32^. Moreover, in other flying insects, this homology even extends to detecting perturbations^81^. Fields dF2 and vF1 in flies are serial homologues of campaniform fields on the radial vein of the wing (dorsal Radius A and ventral Radius A, respectively)^32^. We found that this homology extends to the organization of the motor system for each structure, as the haltere motor system is functionally segregated like the wing steering muscles^47^(Fig. 5). Interestingly, along with their distinct morphologies, the two fields dF1 and vF2 have no clear homolog on the wing. It is possible that the development of these two fields is an evolutionary novelty that–combined with the existing control loop–allowed the haltere to become a multifunctional sensor that provides reafference for controlling flight maneuvers. The breadth of tools available in *Drosophila* combined with classic approaches drawn from biomechanics and biophysics will enable for a fuller appreciation of this enigmatic sensory structure.

## Acknowledgements

We thank Sung Soo Kim for sharing code to track wingstroke amplitude. We used stocks obtained from the Bloomington Drosophila Stock Center (NIH P40OD018537). Gaby Maimon, Michael Dickinson, and Ellie Heckscher also generously shared fly stocks. We thank Marie Suver for helpful comments. Helena Casademunt, Hsin-Yi Hung, Core Park, and Quilee Simeon performed preliminary optogenetic activation experiments at the Kavli Institute for Theoretical Physics at UCSB (NSF PHY-2309135 and Gordon and Betty Moore Foundation Grant No. 2919.02). The data in Figure 5 were collected in the laboratory of M. Dickinson. This work was supported by NINDS-NIH grant 1U01NS131438-01 to J.L.F and B.H.D.; a Searle Scholar Award, a McKnight Scholar Award, NSF grant IOS2221458 to B.H.D; and NSF grant IOS1754412 to J.L.F. The content is solely the responsibility of the authors and does not necessarily represent the official views of the National Institutes of Health.

## Author contributions

Conceptualization, A.V., K.L., J.L.F., and B.H.D.; methodology, A.V., K.L., J.L.F., and B.H.D.; investigation, A.V., K.L., and B.H.D.; resources, J.L.F. and B.H.D.; analysis, A.V., K.L., J.L.F., and B.H.D.; writing, A.V., K.L., J.L.F., and B.H.D.; supervision, J.L.F. and B.H.D.; funding acquisition, J.L.F. and B.H.D.

## Methods

### Lead contact and materials availability

This study did not generate new unique genetic reagents. Further information should be directed to Lead Contact, Bradley Dickerson.

### Animals and tethering

All flies used in this study were 2-to-5-day-old females. For imaging experiments, we raised flies on standard cornmeal medium at 25°C on a 12:12 hour light/dark cycle. For the Chrimson activation experiments, we raised both the parents and offspring in the dark. We added 100µL of 100mM all-trans retinal to the bottles of the parents and 200 µL of the same concentration to the bottles of the progeny.

We expressed GCaMP7f in the haltere afferents by crossing *DB331-GAL4* to w[1118]; P{20XUAS-IVS-jGCaMP7f}su(Hw)attP5; +. We anesthetized the flies at 4°C and tethered them at the anterior notum to a tungsten pin using UV-curing glue. We allowed at least 30 min for recovery before performing experiments.

We expressed GCaMP6f in the haltere steering muscles by crossing w[1118];+; P{y[+t7.7] w[+mC] = GMR22H05-GAL4}attP2 with +[HCS]; P{20XUAS-IVS-GCaMP6f}attP40;+. The haltere muscles are subject to significant motion artifact. Thus, to help stabilize our images, we removed the first two pairs of legs and tethered flies ventrally to a tungsten pin using UV-curing glue placed between the femur of the prothoracic legs and coxae of the mesothoracic legs.

To express Chrimson in the haltere steering muscle motor neurons, we crossed *SS41075-SplitGAL4* with w-; ScO/CyO; UAS-Chrimson.mVenus(w+)/(TM6B,Tb,Hu,e). For the empty vector control, we crossed the empty stable split line w[1118]; P{y[+t7.7] w[+mC]=p65.AD.Uw}attP40; P{y[+t7.7] w[+mC]=GAL4.DBD.Uw}attP2 with w-; ScO/CyO; UAS-Chrimson.mVenus(w+)/(TM6B,Tb,Hu,e). We tethered the flies as in our haltere afferent imaging experiments.

### Flight arena and visual stimuli

For imaging of the haltere campaniform sensilla and steering muscles, we placed flies in the center of a previously described arena^82^ composed of blue light-emitting diodes (LEDs; 470 nm peak wavelength). The arena spanned ± 60° in elevation from the fly’s horizon (32 pixels) and 270° around its azimuth (72 pixels; 3.75/pixel). To accommodate the imaging objective, there was a 90° gap in azimuth on the left side of the arena. We placed one layer of blue filter (Rosco no. 59 indigo) to prevent light from the display from leaking into the camera used for imaging GCaMP activity. Visual stimuli consisted of either wide-field, random dot starfields or a dark stripe that subtended 22.5° on the fly’s retina. Rotational patterns simulated motion at an angular velocity of π rad/s. To test rotational tuning about the yaw-roll and pitch-roll axes, we altered the center of rotation in 30° increments. To test tuning in the yaw-roll plane, we shifted the stimulus from the vertical body axis to the longitudinal axis. To test tuning in the pitch-roll plane, we shifted the stimulus from the longitudinal axis to the transverse body axis. We displayed patterns in random blocks for a duration of 3 s each, five repetitions for each stimulus. To promote flight, we presented flies with a dark stripe on a bright background under closed-loop conditions for 5s between each trial. The pattern then appeared and was still for 1 s before and after each stimulus presentation.

### Flight behavior

To track steering behavior during haltere afferent and muscle imaging experiments, we placed flies within an optoelectronic wingbeat analyzer ^83^. The moving wings cast shadows onto an optical sensor that converts instantaneous wingbeat amplitude into a voltage signal. We acquired wingbeat amplitude data at 2 kHz using a Digidata 1550B or 1440A amplifier (Molecular Devices) for the afferents and muscles, respectively. In our haltere afferent imaging experiments, we also illuminated flies from above with a single IR LED attached to a collimating lens and used a custom MATLAB machine vision script to calculate and record the left-minus-right wingbeat amplitude. In cases where flies stopped flying, we softly blew on them to resume flight.

### Functional imaging

Our method for imaging campaniform sensilla activity on the dorsal side of the left haltere was similar to that described for recording wing muscle activity ^42^. We imaged the haltere campaniform sensilla with a 50x, 0.55 NA objective (Mitutoyo) mounted to 0.75x zoom tube lens on a Thorlabs WFA2001 epifluorescence module, giving us an effective magnification of 37.5x. We collected GCaMP fluorescence using a Retiga R1 camera. The amplifier we used to collect wingbeat amplitude data sent a TTL pulse to an Arduino Due, which triggered the camera at a phase of 0.5 relative to the upstroke of the wings. The Arduino also controlled the excitation LED (M470L3, Thorlabs), which provided 470 nm light to the haltere in three, 1 ms pulses during each exposure of the camera. Images were band-passed filtered by an ET535/50m emission filter (Chroma Technology). We collected TIFF stacks at an exposure time of 22 ms using µManager.

We used a similar method to image the haltere steering muscles. We placed flies beneath a 50x, 0.55 NA objective (Mitutoyo) mounted to a Nikon Eclipse FN1 epifluorescence microscope. In this setup, the fly, arena, and wingbeat analyzer were all mounted sideways to image the muscles. We excited GCaMP6f within the muscles with continuous 470 nm light (M470L3, Thorlabs). We collected images with a QIClick camera (QImaging) after they were band-passed filtered using the same emission filter as in our haltere imaging experiments. We triggered the imaging camera at a phase of 0.75 and used an exposure time of 33 ms.

### Quantification and statistical analysis

We analyzed our imaging and flight behavior data using custom scripts written in Python. For the haltere campaniform imaging experiments, we rigidly registered each image to the mean of all images for the full experiment. We then segmented the mean image of each experiment into two areas of interest, which we interpret as dF1 and dF2, using k-means clustering. After segmenting our images, we computed the change in GCaMP fluorescence F_t_ for each time point. For each campaniform field, we computed the mean baseline fluorescence F_0_ for 0.5 s prior to stimulus motion before computing (F_t_-F_0_)/F_0_, which we term ‘‘ΔF/F’’. We constructed 95% confidence intervals by resampling the population average 1,000 times with replacement from the individual means. To construct tuning curves, we summed each fly’s individual mean fluorescence and wingbeat amplitude signals during the 3 s stimulus period for each stimulus direction.

### Saccade identification

We recorded wing motion and GCaMP activity while flies flew in near-complete darkness to elicit spontaneous turning maneuvers. To record haltere campaniform or muscle activity during spontaneous saccades, we imaged flies while they attempted to fixate a dark, 30° bar under closed-loop conditions.

We identified saccades using a method previously described ^58,84^. Briefly, we first computed the left-minus-right wingstroke amplitude (L-R WBA) and low-pass filtered these data (cutoff frequency of 6 Hz). We then took the derivative of the filtered signal and identified steering events as the local maxima and minima. We defined saccades as the maxima and minima that exceeded both velocity and magnitude thresholds. We then identified the signatures of spontaneous saccades in the haltere campaniform fields and muscles by finding the fluorescence signals that corresponded to a given saccade.

For each saccade, we calculated the mean change in L-R WSA for the 100 ms windows preceding and following each event, triggered at the peak. We then sorted these data to compute decile means. We conditioned our wingbeat amplitude signals by baseline-subtracting the mean of the first 250 ms of a given decile mean. We used this same window preceding the peak in L-R WSA to calculate ΔF/F for the haltere afferent terminals and steering muscles.

### Optogenetic activation of the haltere steering muscles

To track haltere movement, we tethered flies and mounted them within an optoelectronic wingbeat analyzer^83^ with the wingbeat amplitude and frequency sampled at 1 kHz. We mounted a high-speed video camera (Fastec IL5) with a 4x magnification to view the haltere amplitude and frequency, sampling at 2 kHz. We activated the haltere steering motor neurons during tethered flight using a 250 ms pulse of 625 nm light (M625F2, Thorlabs) at a stimulus intensity of 2.8 W/m^2. We then tracked haltere motion using DeepLabCut^85^ and edited with custom MATLAB software. We analyzed all data using custom scripts in MATLAB and Python.

## Data and code availability

The data from this manuscript are published on GitHub at: https://github.com/AnnaVerbe/Flies-actively-tune-the-sensitivity-of-their-multifunctional-gyroscope

**Supplemental Movie 1: *In vivo* imaging of the dorsal haltere campaniform fields during tethered flight.** Top left: Behavioral tracking showing the fly from below and the wingstroke envelope of both wings; left wing amplitude is shown in green. Bottom left: The dorsal campaniform fields clustered by fluorescence activity into regions that correspond with dF1 (red) and dF2 (blue). Right: Extracted signals of the left wingbeat amplitude, fluorescence of the mean area of dF2 and dF1.

**Supplemental Movie 2: Optogenetic activation of haltere steering muscle motor neurons decreases haltere stroke amplitude.** Right: A tethered fly expressing CsCrimson-UAS in the hDVM and hI1 muscles driven by *SS41075-GAL4*, filmed using high speed videography. Traces of haltere motion, tracked using DeepLabCut and custom MATLAB software, are overlaid. Left: The full trace of the haltere envelope (position) before, during, and after optogenetic activation using a 250 ms pulse of 625 nm light. The fly returns to baseline haltere motion slightly after the crimson light turns off.

---

**Figure S1.**
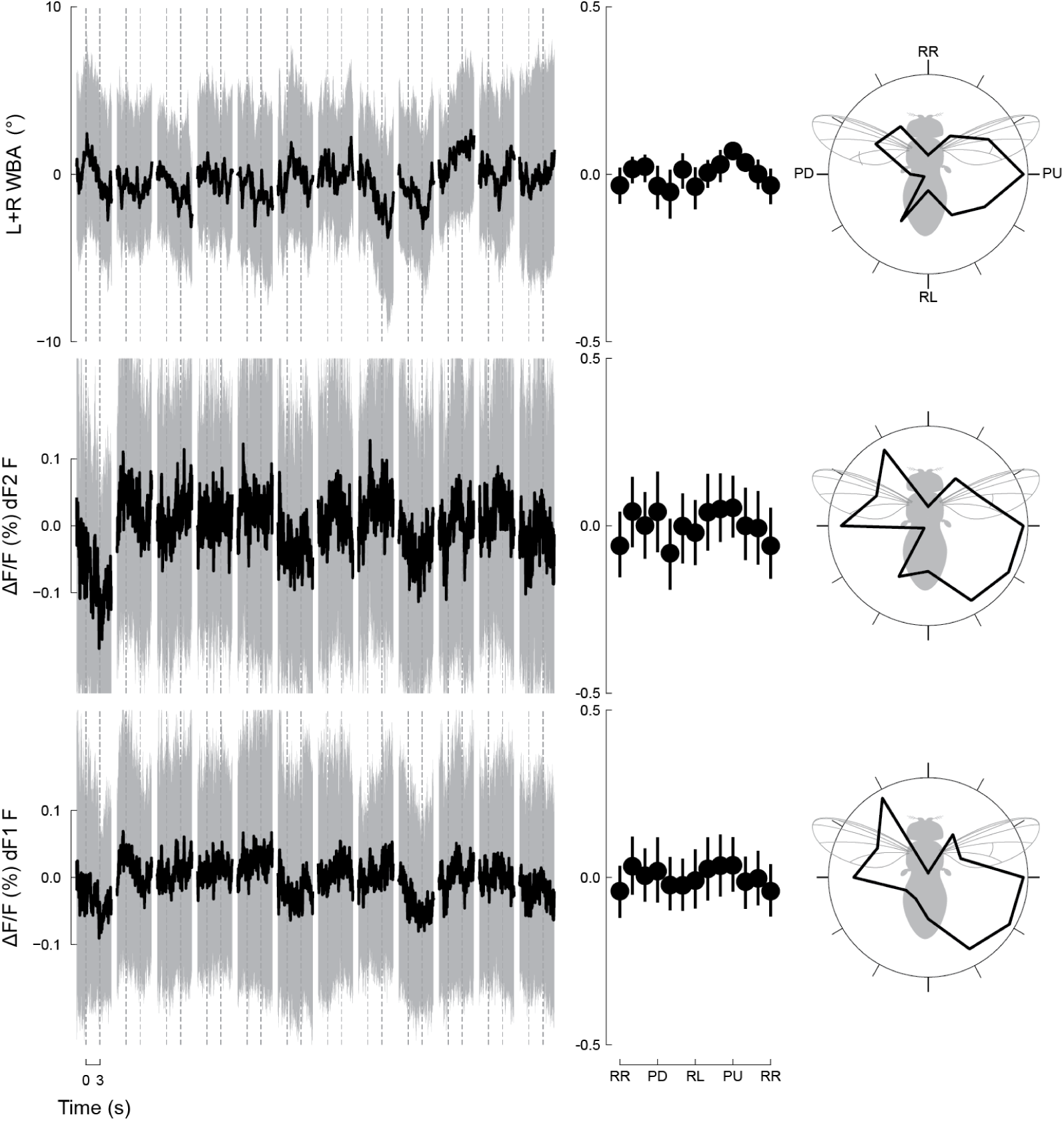
Dorsal campaniform sensilla tuning dynamics about the pitch-roll axis. Related to Figure 2. (A) Averages of wing kinematics and dF1 and dF2 signals for the 12 types of visual motion stimuli around the pitch-roll axis: pitch down (PD), pitch up (PU), roll right (RR), and roll left (RL). Envelopes represent ±95% CI. (B) Stroke amplitude and halteres campaniform sensilla activity plotted as functions of rotational orientation in the midsagittal plane. The curves were constructed by determining the average signal over the 3 s following stimulus onset. (C) Mean tuning functions from (B) plotted in polar coordinates. n = 12

**Figure S2.**
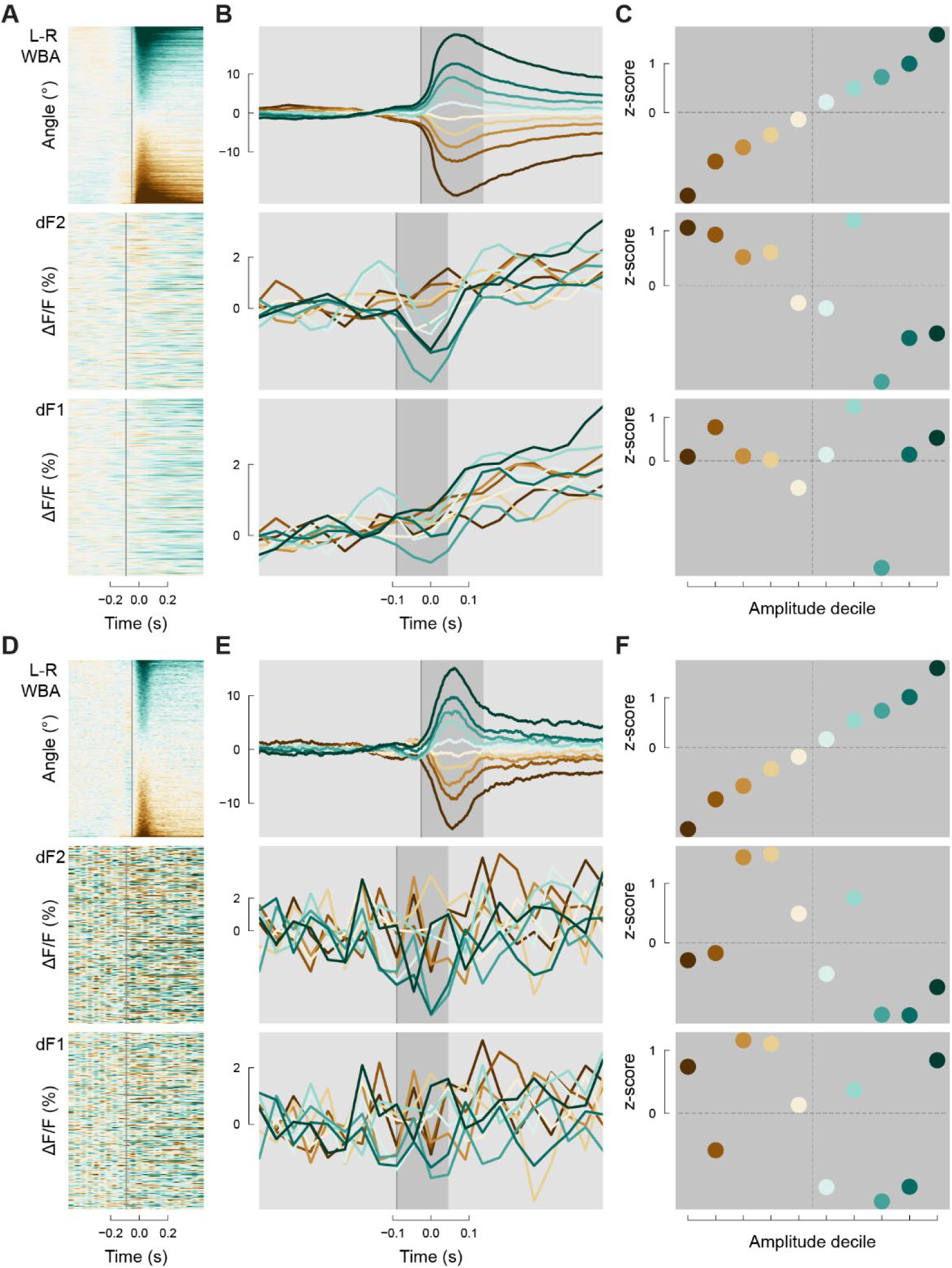
Spontaneous saccades during presentation of a static visual pattern. Related to Figure 4. (A) Raster plots of all saccades sorted by the change in left-minus-right wingbeat amplitude. (B) Saccade-triggered averages of wing kinematics and dorsal halteres signals sorted into amplitude deciles, ranging from the largest rightward turns (0^th^–10^th^ percentile, green) to the largest leftward turns (90^th^–100^th^ percentile, brown). (C) Plots showing mean response for each decile during the saccade (dark grey bar in B). n = 6748 saccades from 19 flies. (D-F) Same as A-C, but for control experiments expressing GFP in the haltere campaniforms where flies tracked a dark stripe in closed-loop. n = 4521 saccades from 6 flies.

